# Drift correction in localization microscopy using entropy minimization

**DOI:** 10.1101/2021.03.30.437682

**Authors:** Jelmer Cnossen, Tao Ju Cui, Chirlmin Joo, Carlas Smith

## Abstract

Localization microscopy offers resolutions down to a single nanometer, but currently requires additional dedicated hardware or fiducial markers to reduce resolution loss from drift of the sample. Drift estimation without fiducial markers is typically implemented using redundant cross correlation (RCC). We show that RCC has sub-optimal precision and bias, which leaves room for improvement. Here, we minimize a bound on the entropy of the obtained localizations to efficiently compute a precise drift estimate. Within practical compute-time constraints, simulations show a 5x improvement in drift estimation precision over the widely used RCC algorithm. The algorithm operates directly on fluorophore localizations and is tested on simulated and experimental datasets in 2D and 3D. An open source implementation is provided, implemented in Python and C++, and can utilize a GPU if available.

## 1. Introduction

Single molecule localization microscopy (SMLM) has provided unique insights into molecular biology by allowing optical microscopy to be used beyond the diffraction limit (∼ 250 nm). It is possible to separate the signal of individual fluorescent molecules, by using fluorescent molecules that can individually turn on and off. This way their location is estimated with high precision (∼ 10*nm*). Many images of the same sample are recorded, processed, and combined into a single super-resolution image.

Slight drift of the sample during acquisition results in a loss of resolution. Therefore, any super-resolution reconstructions need to be compensated for drift during the measurement. This can either be done through active compensation during the measurement or in post-processing. Early work on active drift correction using a fiducial spot created with a pinhole achieved sub-nanometer localization precision [1]. Other methods use fiducial markers embedded in the sample [2, 3], or use image processing of bright field imaging from the sample itself to estimate drift [4]. If active drift correction is limited to purely z-drift (along the optical axis), commercially available interferometry autofocusing systems are frequently used. This still leaves 2D drift uncorrected, but because drift within the focal plane does not deteriorate the image quality, it can be corrected afterwards without any negative consequences as long as the estimation of the drift is accurate enough. If the sample is not sufficiently static, fiducial markers can be embedded in the sample [5]. For SMLM however, although different sets of fluorophores are visible through time, the underlying structures that are being imaged are often static and thus suitable to use as input for a drift estimation algorithm. Several such approaches have been explored: [6] estimated drift by binning localizations over time, and performed image cross-correlation of the first bin against all remaining bins. Redundant cross correlation [7] (RCC) improves on this by computing the cross-correlations between all bin pairs. Due to the quadratic scaling of the number of bin-pairs, this method is fast if the number of bins remains relatively small (<20), but can be much slower if a higher time resolution is required. RCC was further extended into 3D SMLM in ZOLA-3D [8].

To our knowledge, further improvements of post-process drift correction over RCC have been limited: [9] demonstrated Bayesian sample drift inference (BaSDI), an optimal approach using Bayesian statistics and expectation-maximization, in which the drift can be estimated per-frame. However, this approach internally uses an image-based representation of the localizations, which means that memory use and compute time do not scale well with either large fields of view, or very precise measurements. Sub-nm precise drift estimation has been demonstrated [10] specifically on DNA-PAINT origami, using markers and taking advantage of the separable clusters and known sparsity of binding sites, but rendering the approach incompatible with larger and denser biological structures such as microtubules. In practice, we note that in the absence of fiducial markers or active drift correction, RCC is currently the method of choice for researchers using SMLM. Despite its widespread use, the precision and bias of RCC and how much room is still left for improvement, is poorly understood.

Here, we propose a new method for drift estimation in both 2D and 3D datasets based on the minimization of a bound of the entropy. Our method, Drift at Minimum Entropy (DME), runs directly on localization data, supports both 2D and 3D drift estimation, and does so with a compute time similar to RCC. The method is tested on DNA-PAINT origami measurements (Figure 2), simulated SMLM measurements of filaments Figure 4, experimental STORM imaging of microtubules (Figure 5) and simulated 3D microtubules (Figure 6). The outline of the paper is as follows: in section 2.1 to section 2.4, we describe the concept and mathematical definition of the estimation method; in section 2.5 describes preparation and measurement of a DNA-origami sample that was used to compare fiducial markers and localization based drift estimation; section 3 describes the various experiments we performed, comparing the performance of DME, RCC and fiducial markers drift estimation; and section 4 discusses the results and open questions.

## 2. Methods

### 2.1. Drift at Minimum Entropy

The true emitter position has, under mild assumptions, a normal distributed error with respect to its true position [11]. The localization microscopy reconstruction can therefore be expressed as a probability distribution

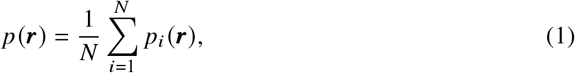

where *N* is the total number of localizations and ***r*** is a *K*-dimensional position vector representing the emitter position. *p*_*i*_ (***r***) is the probability distribution of a single localization:

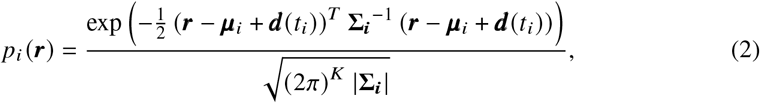

with ***µ***_*i*_ as the estimated position, **Σ**_***i***_ the estimated covariance matrix, and ***d*** (*t*_*i*_) is the drift in frame *t* of the *i*-th emitter.

The drift is estimated by minimizing an upper bound on the statistical entropy of *p* (***r***). We note that *p* (***r***) is a Gaussian Mixture Model (GMM) with equal weights for each component [12]. The entropy of a GMM does not have a closed form solution, but various closed form upper bounds have been found [13–15]. DME uses a variational approximation of the entropy from [15] for its computational simplicity:

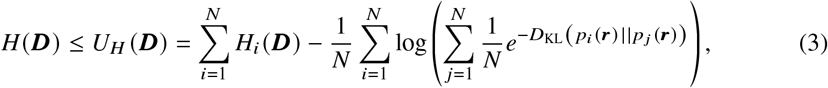

where *H* (***D***) is the entropy of distribution *p* (***r***), *H*_*i*_ (***D***) is the entropy of a single GMM component *p*_*i*_ (***r***). The entropy of a single GMM component is given by 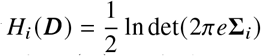, which will stay constant under drift change. Therefore, the first term of *U*_*H*_ (***D*)**^**2**^ can be ignored during the drift estimation. *D*_KL_ (*p*_*i*_ (***r***) || *p* _*j*_ (***r***)) is the Kullback-Leibler divergence between the probability distributions for localization *i* and *j*. In our implementation, we assume localization errors to be uncorrelated between axes, so the non-diagonal terms of ∑ are zero. *D*_KL_ (*i, j*) can then be expressed as follows:

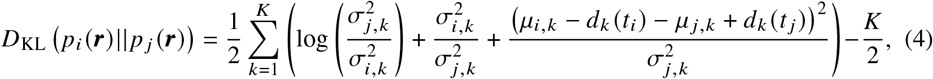

Here 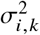 is the *k*-th diagonal of the covariance matrix ∑_*i*_ for localization *i*. To compute the covariance matrix we use the Cramer Rao Lower Bound (CRLB), which defines a lower bound on localization precision of a fluorophore [16]. While Equation 3 appears to be very computationally demanding, scaling quadratically with the number of localizations, this is effectively not the case in typical SMLM datasets. The exponential term 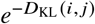 goes to zero with increasing distance between localizations. To decide which localizations to include in the inner sum of Equation 3, we set a fixed distance threshold based on the average CRLB times 3 and ignore any localizations further than that. At that point, assuming equal CRLB, the entropy contribution of a localization pair is at the 3*σ* level (0.3%). An efficient Kd-tree [17] implementation is used to find localizations within this distance. Whenever the drift estimate has moved more than half the distance threshold, the localization neighbors are recomputed.

Runtimes of DME are on the same order of magnitude as those of RCC, typically less than a minute (see figure Figure 2,b). For large datasets, if compute time is still too long or too much memory is used, the number of localizations can be reduced to include only the brightest spots. The default maximum is set to 1M localizations, which we found to be more than sufficient for drift estimation in all tested datasets.

### 2.2. Cubic spline based drift parameterization

To reduce the number of estimated parameters, we define the drift ***d*** (*t*) as a Catmull-Rom spline i.e. a piecewise cubic spline where the value is determined by its 4 neighboring nodes. Such a spline function is computationally efficient to evaluate and differentiate, and tunable to the specific dataset by changing the number of knots. To do this, the global frame index t has to be translated into a local coordinate on the spline *τ* and an index *j*, defining which of the spline segments corresponds to frame *t*. The drift at frame *t* is defined as:

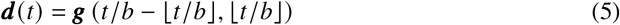

where

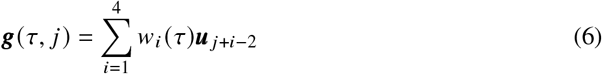

with

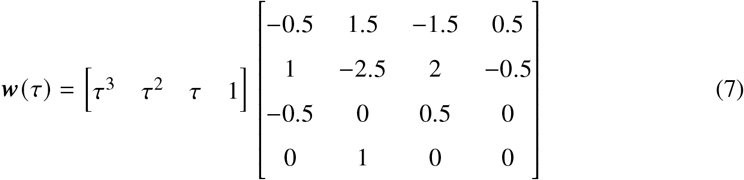

here *j* is the spline segment index, *τ* is an interpolating fraction between 0 and 1, **u** _*j*_ is the *j* -th spline control point, and *w*_*i*_ (*τ*) the *i*-th element of ***w*** (*τ*) the Catmull-Rom spline. We define *M* drift spline control points, *u* _*j*_ with *j* ∈ {0, *M* − 1} and *M* = [*L b*], with *b* being the binning size, i.e. the number of frames in a single spline segment and *L* being the total number of frames.

### 2.3. Gradient descent

The drift is estimated by finding a set of values for ***u***_*k*_ that minimize *H* (***D*** (***u***_*k*_)) (Equation 3), using gradient descent (see supplementary note Y). Figure 1 illustrates the general idea of the algorithm. By dynamically adjusting the gradient step size we optimize the speed of convergence. Because DME uses gradient descent in a complex optimization landscape, it is susceptible to converge to a local minima. To prevent this, we use 2D RCC with 10 bins to quickly compute a rough initial estimate. In 3D drift estimation, we also run a coarse optimization step after the RCC. In the coarse optimization, the optimization landscape is smoothed by doubling the *∑*_*i*_ values of all localizations used during drift estimation.

**Fig. 1.**
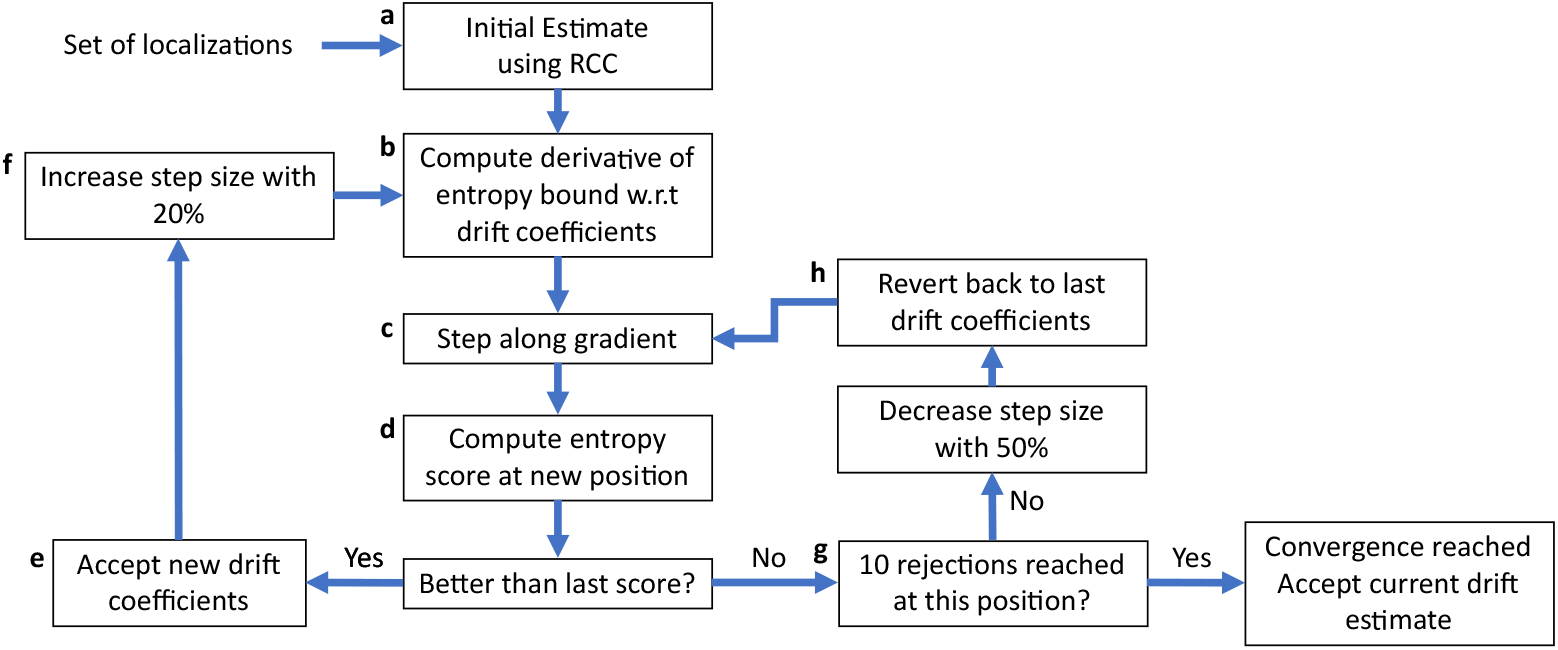
Schematic representation of the DME algorithm. **a**) The RCC algorithm is used in X,Y with a minimal number of bins to create an initial estimate of drift. This makes DME more robust in cases of large drift displacement, and can alternatively be replaced with DME running on a large bin size. **b**). Given the spline-based definition of drift, we compute the derivative of the entropy upper bound w.r.t all of the spline coefficients that describe the drift. **c**) Using the computed gradient and current step size, we compute a new set of spline coefficients. **d**) The entropy score is defined as upper bound of the entropy, except for the parts that stay constant during change of the drift. If the entropy score has improved, we accept it as a new drift estimate (**e**). **f**) Gradient descent can converge very slow if a sub-optimal step size is used. To maintain a close to optimal step size, we increase it on successful steps and decrease it whenever the entropy score is not improving. **g**) The algorithm is stopped when there are no improvements in the score, even after decreasing the step size 10 times. At that point, we can conclude we have reached a (global or local) minimum in the entropy score.

### 2.4. Spot detection and localization

All conversion of raw images to 2D and 3D localizations has been done using a custom implementation of conventional SMLM methods. Details can be found in the supplement.

### 2.5. DNA origami experiments

#### 2.5.1. Sample preparation

The DNA origami plate (Tilibit nanosystems, Munich, Germany) was designed using CADNano [18] on a square lattice design as described previously [19]. Assembly occurred in a 100 µL reaction volume containing 10 nM p8064 scaffold strand, 100 nM core staple strands, 100 nM target handles, 100 nM biotin handles in 1x TE folding buffer with 11 mM MgCl2. The DNA plate was annealed by setting the thermocycler to 65°C for 10 min, after which a temperature gradient was applied from 60°C to 40°C in steps of 1°C/hour. Subsequently, the assembled plate was separated from the staple, target and biotin handles using an Amicon spin filter (100 kDa). The plates were kept inside T50 buffer (50mM Tris-HCl, pH 8.0, 50 mM NaCl) supplemented with 11 mM MgCl2 (origami buffer).

Quartz slides were pegylated according to Chandradoss et al. [20] to avoid non-specific interactions. After assembly of microfluidic plates, slides were incubated with streptavidin (0.1 mg/mL, ThermoFisher) for 1 min. followed by washing with origami buffer. DNA origami plates were introduced inside the chamber at a concentration of approx. 50 pM and incubated for 1 min after which the chamber was rinsed with origami buffer. DNA imager strand was subsequently introduced into the chamber with imaging buffer (50 mM Tris-HCl, pH 8.0, 50 mM NaCl, 1 mM MnCl2, 5 mM MgCl2, 0.8% glucose, 0.5 mg/mL glucose oxidase (Sigma), 85 ug/mL catalase (Merck) and 1 mM Trolox (Sigma)).

#### 2.5.2. Microscope setup and data acquisition

Experiments were performed on a custom built prism-TIRF setup. An inverted microscope (IX-73, Olympus) is used with a 532 nm (Sapphire 532-100 mW CW CDRH) laser to excite the DNA imager strand. The resulting fluorescence signal was collected using a 60x water immersion objective (UPLSAPO60XW, Olympus). A longpass filter blocks the excitation light (LDP01 - 532RU - 25, Semroc) after which the fluorescence signal is then projected onto the EM-CCD camera (iXon3, DU - 897 - C00 - #BV, Andor Technology). The images were recorded using Andor Solis.

## 3. Results

We quantified the performance of DME in a range of experimental conditions and simulations. An experimental comparison was performed on a DNA-PAINT sample that contains both DNA origami nanostructures and fiducial markers (Figure 2). The recorded images were processed in our localization pipeline (see supplement section 3), and Picasso Render [21] was used to manually mark all the beads to separate them from the DNA-paint localizations. 18 beads were removed from the set of localizations, and 4 best beads (with the least noisy traces) were used for fiducial marker based drift correction. The non-bead localizations from this experiment were drift-corrected using using RCC, BaSDI and DME at different frame bin sizes. It can be seen that DME is able to reach an equal or better precision as fiducial markers at small bin sizes. BaSDI is also able to do so, but at a much longer compute time and adds discretization error due to discrete grid in which the drift is represented. The estimated precision of localization clusters agrees with the CRLB (and is even slightly lower, potentially due to camera calibration of photon counts). This indicates that for DME, BaSDI, and fiducial markers, the drift is fully corrected and the remaining cluster width can be attributed to localization error. In this particular DNA-PAINT acquisition, RCC will make cluster precision a few nanometer worse, and can correct drift only at a time resolution that is orders of magnitude worse than DME. A comparison of localization-estimated drift with fluorescent bead drift traces in Figure 2.**d** shows a smoother error distribution for DME compared to BaSDI, likely caused by the discretization error of BaSDI. As it is hard to estimate true drift estimation precision without the ground truth drift, we performed an additional simulation (Figure 3) with clusters of localizations (similar to DNA-PAINT measurements). Here, it is clear that DME has a better precision versus compute time ratio, and outperforms BaSDI and RCC on a typical SMLM localization reconstruction.

**Fig. 2.**
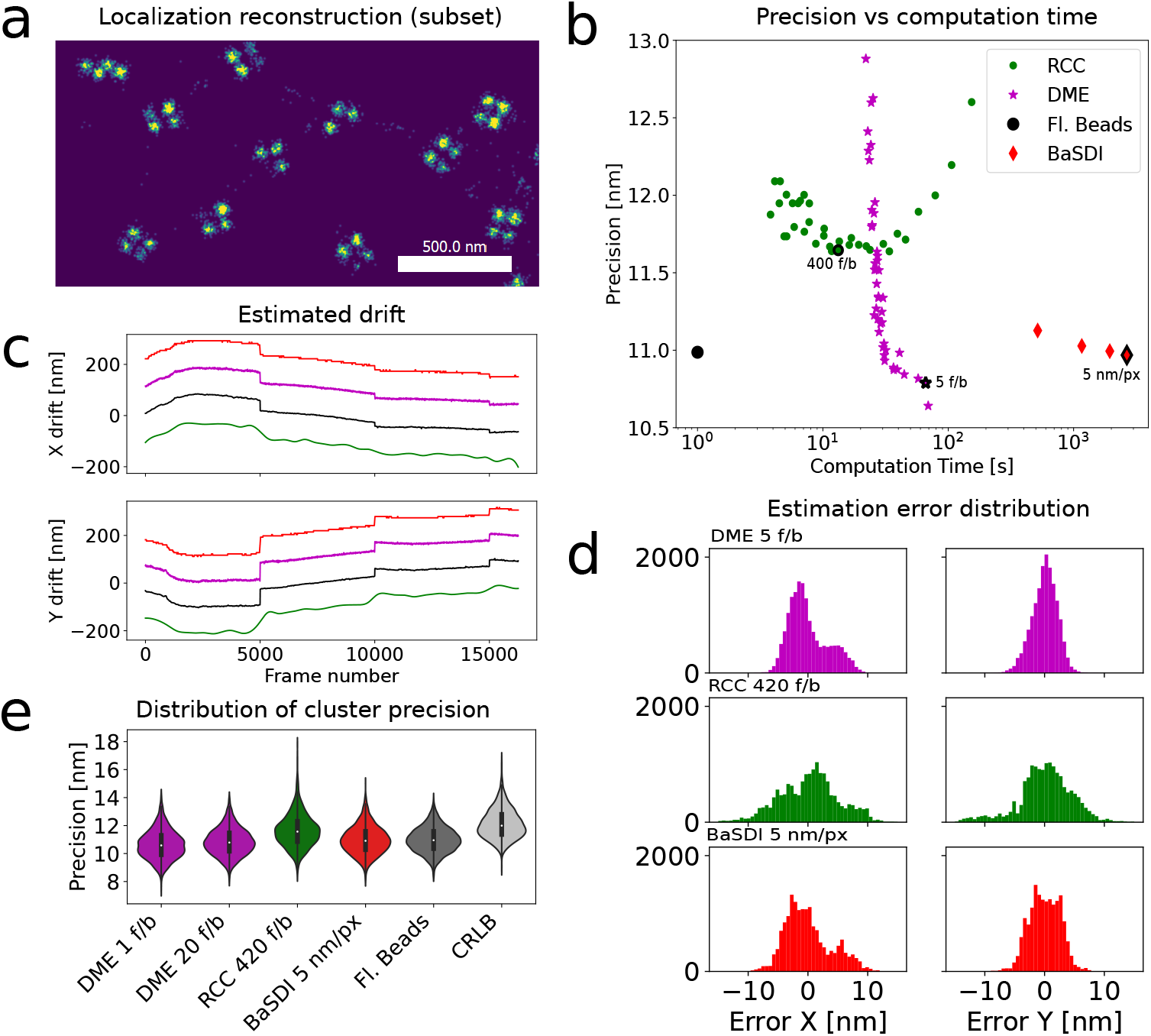
Drift correction using 3 different methods for a DNA-PAINT experiment done on a TIRF microscope, imaging 16256 frames at 200 ms exposure time. The sample consists of square DNA origami binding sites of 60 nm x 80 nm, with fluorescent tetraspeck beads as fiducial markers. A subset of the binding sites are shown in **a**, visualized using Picasso Render [21]. Converting the image data to localizations was done as described in methods. The DNA-PAINT localizations have an average emitter photon count of 462 photons, with 4.4 photons/pixel background, whereas the bead localizations have an average photon count of 12635 photons, and 28.7 photons/pixel background. The disparity in background is most likely caused by PSF model mismatch with the 2D Gaussian PSF. **b**) compares drift estimation precision with required computation time. The drift estimate from fiducial markers (black) is compared to DME (purple), RCC (green) and BaSDI (red) on the localizations without beads. The dataset was chosen as it has several discrete jumps in it due to pausing of the recording, which is suitable to demonstrate the improved time resolution of the DME algorithm. The precision is estimated from the standard deviation of localization positions corresponding to a binding site (cluster), with the mean standard deviation of the clusters plotted. The plotted drift traces (**c**) are shifted by one pixel each to prevent overlap in the plot. Frame bin sizes were chosen to be optimal for each algorithm, 60 frames/bin was used for DME and 400 frames/bin for RCC (also labelled in **b**). For BaSDI, a subset of the dataset was used, cropped to 120× 120 pixels, and the number of frames was reduced by 4x using binning. 5.414 nm/pixel was used for as BaSDI grid spacing. The difference between the estimate from fiducials markers and the estimate from the algorithms (**d**) indicates potential drift bias on the shape of resolved structures in a sample. Noticeable is that in the X axis the DME drift difference does not follow a Normal distribution. Since this shape is also seen in the BaSDI drift, it could be caused by a bias in the fiducial marker localization due to PSF abberrations. Panel **e** shows standard deviation of localizations in 2604 clusters. Clusters with < 30 localizations are discarded.

**Fig. 3.**
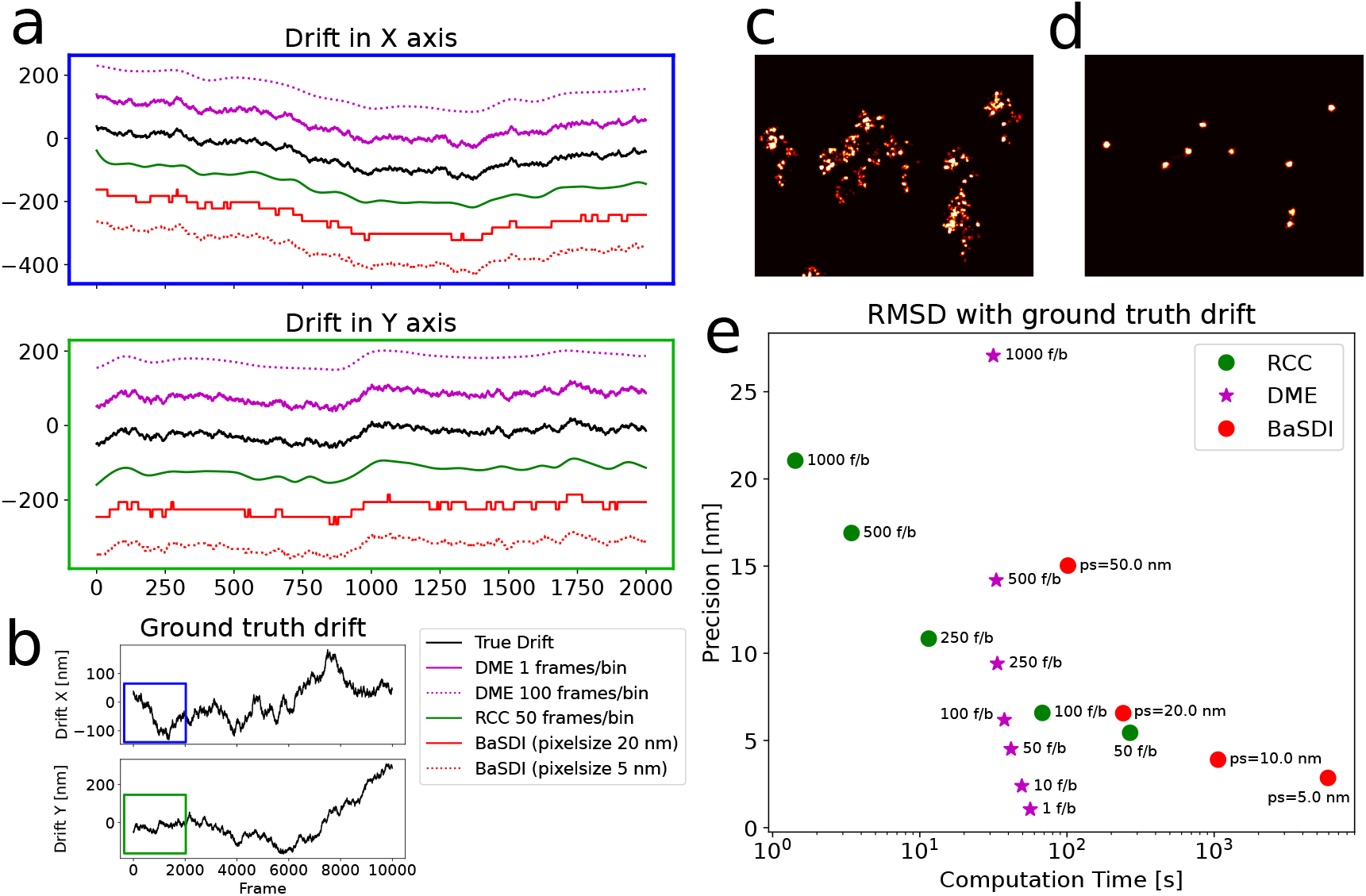
Drift correction using various algorithms on a simulation of blinking spots at fixed positions, similar to DNA-PAINT measurements. The drift (**b**) was simulated using a cumulative normal distribution, sampling a new offset from a 𝒩 (*µ* = 0.02nm, *σ*^2^ = 4nm^2^) A sub-interval of the drift is shown in (**a**). 1000 binding sites were simulated within an area of 300×300 pixels (100 nm/px), with 3000 photons per localization and 10 photons/pixel background. 10000 frames were simulated resulting in a total of 353901 localizations. A subsection of the localization dataset is shown before drift correction (**c**), and after drift correction (**d**). The root-mean-square deviation w.r.t the ground truth drift is plotted versus the required computation time (**e**). Picasso RCC with bins smaller than 50 frames fails to complete due to too few localizations per bin. It can be seen that with this particular set of parameters, such as image size, localization precision and underlying structure, DME is computationally more efficient than BaSDI and achieves a better estimation precision. The best precision for DME, RCC and BaSDI plotted here are 1.05 nm, 5.44 nm and 2.85 nm resp.

DNA origami samples as demonstrated in Figure 2 have a relatively large number of localization events per binding site. Real biological structures typically use a similar number of frames in the acquisition, but the localizations from these frames are spread over a many more fluorescent molecules in the sample. We therefore tested the algorithm performance on simulated microtubule acquisitions, shown in Figure 4.

**Fig. 4.**
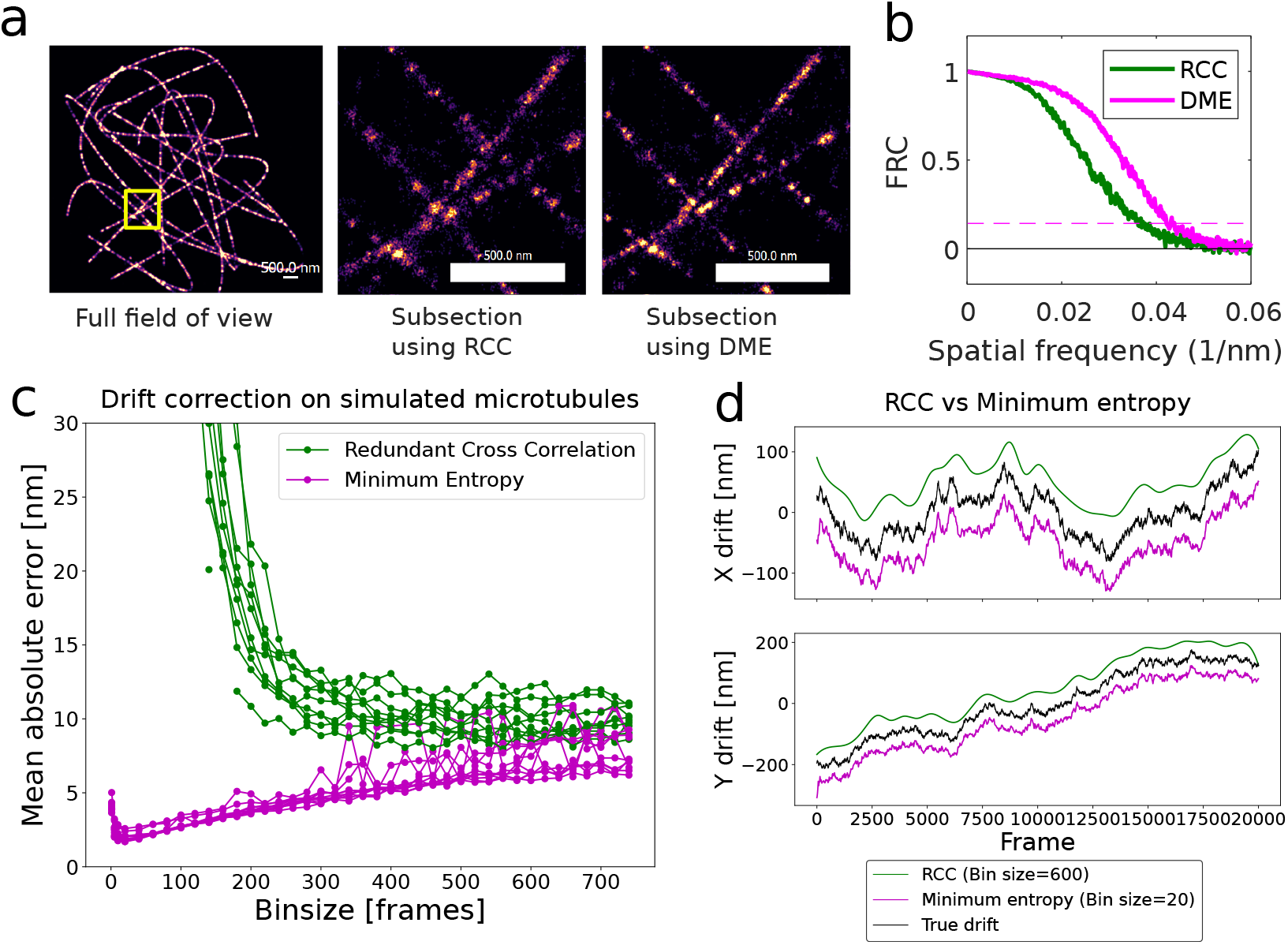
Drift correction on simulated microtubule measurements. To estimate the true error of drift correction methods, we generated 10 datasets with binding sites positioned along randomly generated 2D splines (see panel **a**). Each dataset has 20000 frames of 100×100 pixels (100 nm/pixel), each sampled from Poisson distributed noise. The simulation was done with 20 photons/pixel of fluorescence background, between 500 and 1500 photons per emitter per on frame, and an average of 10 emitters per frame with an average on-time of 10 frames. Drift was simulated as cumulative normally distributed noise, typically around a few pixels as is common in real measurements. (**c**) For each dataset, both RCC and DME drift correction was performed for a range of binning sizes. The mean absolute error (MAE) in the estimate is plotted, with each line representing one of the 10 datasets. (**d**) The true drift trace plotted next to both estimates. Bin sizes are chosen to be optimal for each the algorithm in terms of MAE (20 fr/bin for DME, 600 fr/bin for RCC). (**b**) The FRC for both RCC and DME is plotted for the same dataset as (b), showing a resolution improvement from 27.15 nm (RCC) to 23.35 nm (DME). (**a**) The dataset used in (**b**) visualized, with a subsection shown in yellow indicating the zoomed in area.

It can be seen that DME outperforms RCC on (sparser) microtubule localization microscopy data: when the number of localizations within one bin becomes a limiting factor for RCC, DME will still perform well. The effect of improved drift correction on this particular dataset can be clearly seen in the two zoom-in sub panels. The visual improvement can also be partly due to the shape of the drift trace, as the image blurring might be more significant for the simulated cumulative Gaussian noise in Figure 4 than for the thermal drift seen in Figure 2.

To quantify DME performance on experimental microtubule data, we used a publically available STORM microtubule dataset from SIMFLUX [22]. Results of this are shown in Figure 5. The dataset is split up in two subsets with equal number of localizations. The drift is estimated independently on each of these subsets, which allows us to compute an estimate of the drift estimation precision. The histograms of the difference between these two drift traces for RCC show a significantly non-Gaussian shape of errors, which could indicate potential artifacts of several nanometers in size. This non-Gaussian distribution is caused by the combination of having few time points in the RCC result, and the spline interpolation to interpolate between those points. In typical STORM measurements with a localization precision of >10 nm these artifacts will be hidden, but they could become apparent for more precise localization microscopy methods.

**Fig. 5.**
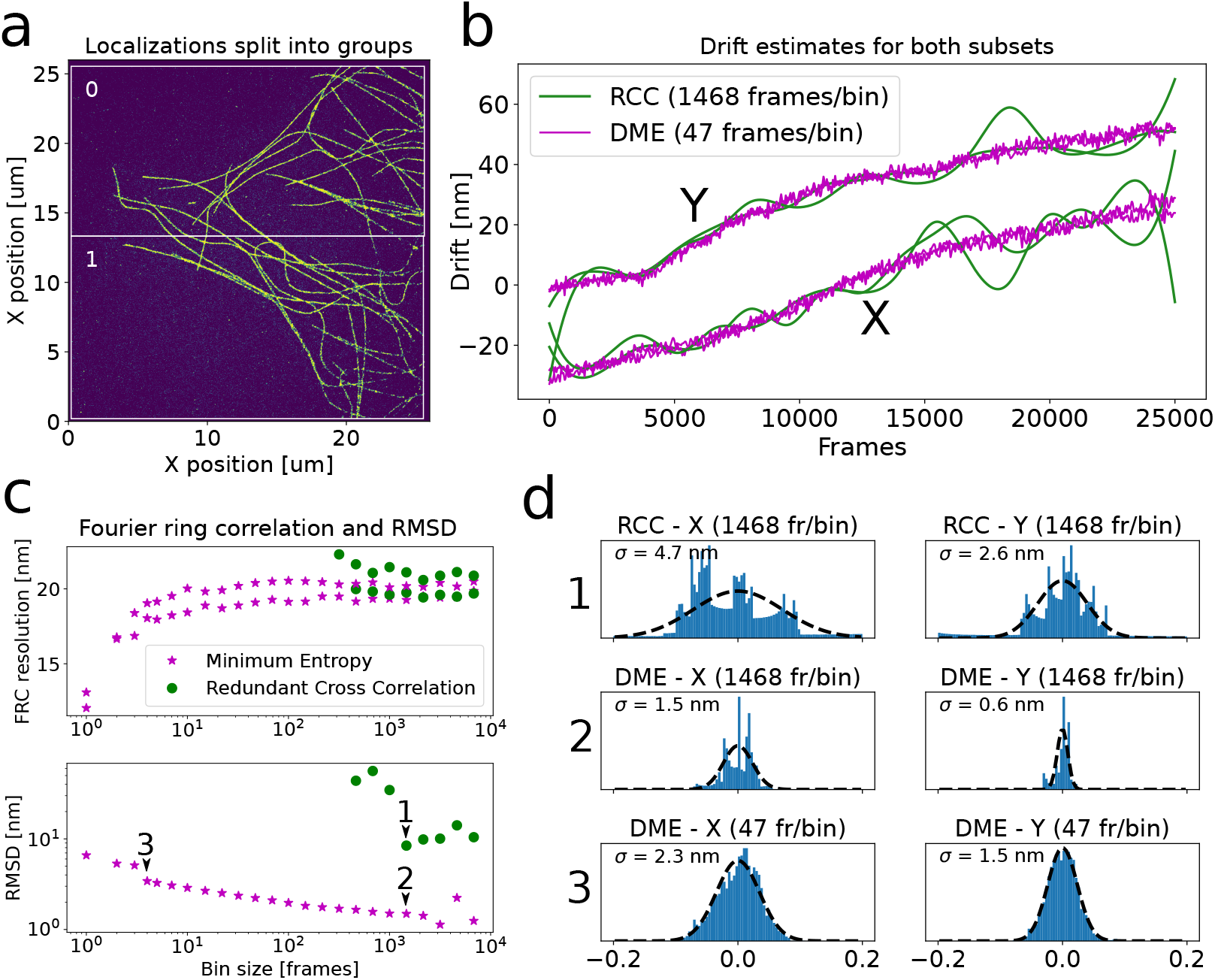
Drift correction comparison done on a STORM acquisition of microtubules from SIMFLUX [22]. The comparison was done on uniform excitation localizations, by summing every 6 frames to get raw images equivalent to uniform excitation measurements. The resulting localization dataset was split up in two sets of 410401 localizations each (**a**), and both RCC and DME was performed on each subset using different binning sizes (**b**). The Fourier ring correlation (FRC [23]) of both subsets, and root mean square difference (RMSD) between the two drift estimates is plotted for a range of drift estimation bin sizes (**c**). (**d**) Histograms of the difference between the two drift estimates, for the two methods in both axes. Apparent is that DME errors are normally distributed at small binning sizes (**d.3**), or have a low variance compared to RCC at equal binning size (**d.1** and **d.2**). RCC is likely creating drift related artifacts on the scale of ∼ 2 nm in the result due to errors not being normally distributed.

The computed Fourier ring correlations (FRC) show a surprising drop in FRC at low bin sizes. After visual inspection of microtubule structures, no clear improvement can be seen. From this we conclude that the FRC improvement is at least partially due to overfitting. At a binsize of 1 frame, each frame on average has only around 16 localizations. At such low counts, we argue that the DME algorithm will move localizations directly on top of each other and assign a non-physical drift that coincides with an FRC improvement. The increase in drift estimation precision as measured by the RMSD (Figure 5, panel C) around low binsizes also supports this conclusion.

The application of DME on 3D drift estimation is demonstrated in a simulated experiment, shown in Figure 6, where two experimentally measured PSFs were used to generate 2 3D microtubule datasets. In 3D drift estimation, we again computed an initial estimate in 2D using Picasso’s drift estimation implementation. Then, we applied a coarse drift estimation by running DME with a larger CRLB assigned to the localizations, which smoothes the optimization landscape and prevents the Z drift estimation getting stuck in a local minima. One caveat of this approach is that the number of neighboring localizations increases quickly with an increasing CRLB, so it quickly increases compute time and memory use. This could alternatively be replaced with a 3D estimate using RCC as done in ZOLA-3D.

**Fig. 6.**
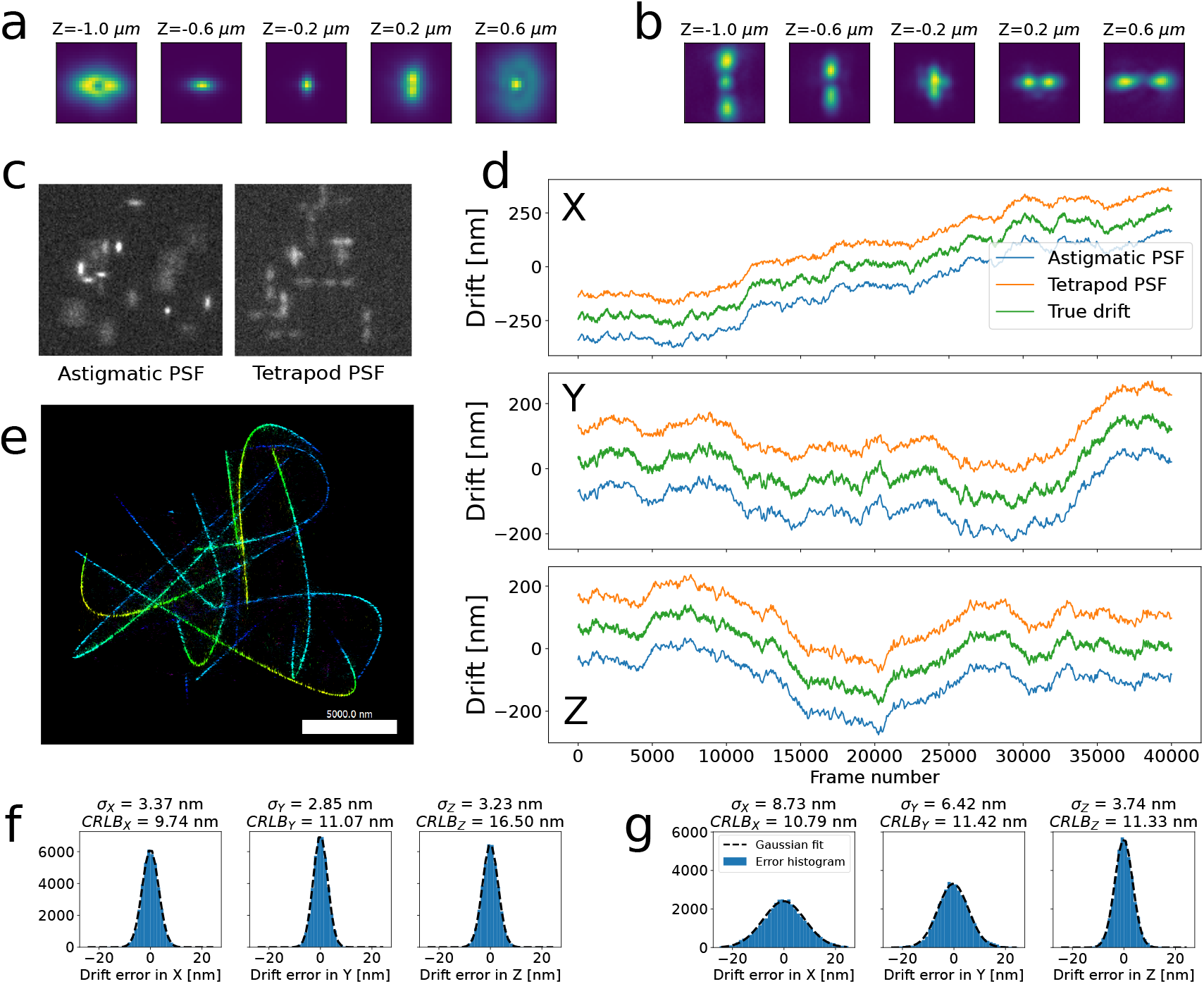
3D drift correction on 2 simulated SMLM measurements of microtubules, done the same as in Figure 4 but spread over a 2 *µm* Z range. The simulation was done with 10 photons/pixel background, and an average of 3000 photons/emitter. Both datasets use the same ground truth structure and drift, but two different experimentally measured PSFs were used: an astigmatic PSF (Tubulin-A647-3D [24]) (**a**), and a tetrapod PSF from our custom build microscope using a deformable mirror (**b**). (**c**) An example of raw frames from the 2 datasets. The simulated camera frames were processed by our 3D localization pipeline as described in the supplement 3.C. (**d**) The drift was estimated first using RCC, followed by a coarse estimation using 4x larger localization ∑ values and 2000 frames per bin, and finally a precise estimation using 50 frames per bin. (**e**) 3D rendered version of the tetrapod PSF dataset, using Picasso Render [21] and Z position encoded as color. (**f** & **g**) The drift estimation errors plotted in histograms for both astigmatic (**f**) and tetrapod (**g**) datasets, showing a clear normal distribution for the drift estimation errors. *σ*_*x, y,z*_ and *CRLB*_*x, y,z*_ indicate the standard deviation of drift estimation error and mean CRLB of invidiual localizations, resp. The difference in drift error can be attributed to different numbers of localizations in the datasets. Due to difference in size and shape of the PSF, spot detection efficiency varies between the datasets and the astigmatic dataset has 241910 localizations whereas the tetrapod data has 144443.

## 4. Discussion and conclusion

On experimental DNA-PAINT origami measurements, DME shows an approximate five fold improvement over RCC in drift estimation precision on DNA-PAINT nanorulers, and is able to achieve the same localization precision as fiducial marker based drift correction (Figure 2.b). We show that DME achieves a better drift estimation precision and is able to do so at a much smaller frame binning size (Figure 5.d), indicating a higher time resolution. In a typical SMLM reconstruction, a binning size of around 50 frames per bin is optimal for DME, versus a 500 for RCC (Figure 4.c). Depending on the drift in the system, this can result in an improvement in image quality as measured by the FRC (Figure 4.b) and localization cluster precision (Figure 2.b). Simulated SMLM acquisitions demonstrate that DME is able to perform precise 3D drift estimation as well, at a near-isotropic 3D precision (Figure 6, panel f and g). Compared to the BaSDI drift estimation algorithm, DME achieves a similar or better drift estimate with a much lower compute time (Figure 2.b). We find that performance of the drift estimation depends very much on the characteristics of the datasets. While BaSDI performs very well on small, sparse datasets, we found it to be prohibitively slow on realistic larger datasets.

The improved performance of DME on sparser datasets also makes it feasible to estimate the precision of the drift estimation itself by cross validation (Figure 5.c). Our implementation reports this using the RMSD between the drift estimates of the dataset split two-fold, which is used to inspect the quality of the DME output and tune the bin size. DME utilizes a gradient descent approach which requires a reasonable initial drift estimate. RCC gives a robust initial estimate, but alternative solutions are also possible. For example, stochastic gradient descent as frequently used in deep learning might improve DME convergence. Additionally, the minimum entropy metric can potentially be used for other SMLM applications, such as alignment of different sample FOVs in image stitching, or multicolor channel alignment.

In conclusion, we developed a novel algorithm for drift estimation on point-based datasets using a minimum entropy optimization metric. We anticipate that for many experiments, especially those with sparse binding sites as found in Figure 2, DME can replace fiducial markers and achieve the same image resolution, simplifying sample preparation. The method scales well to 3D drift estimation, and with 3D SMLM becoming more popular and mainstream, we expect this algorithm to find widespread practical use.

## 5. Code and data availability

Code and example data is available is freely available at the link below. Original image data is available on request. https://github.com/qnano/drift-estimation

## Acknowledgments

J.C. designed the algorithm, performed measurements and simulations. T.J.C. created the combined fiducial marker + DNA-PAINT samples and performed measurements. J.C., T.J.C and C.S wrote the paper with input from C.J.

## Disclosures

The authors declare no conflicts of interest.

See Supplement 1 for supporting content

