## Supplementary Information for "Drift correction in localization microscopy using entropy minimization"

### Supplemental information: Drift correction in localization microscopy using entropy minimization

#### 1. GRADIENT DESCENT ON THE MINIMUM ENTROPY OPTIMIZATION METRIC

As described in the main text, DME uses the following upper bound on the entropy from [1] as optimization goal in a gradient descent approach:

$$U_H(\boldsymbol{D}) = \sum_i^N H_i(\boldsymbol{D}) - \frac{1}{N}\sum_i^N \log\left(\sum_j^N \frac{1}{N} e^{-D_{\text{KL}}(i,j)}\right), \tag{S1}$$

where $\boldsymbol{D} = \{\boldsymbol{d}(1), \cdots, \boldsymbol{d}(L)\}$ defines the drift at every frame.

$D_{\text{KL}}(i,j)$ is short for $D_{\text{KL}}\left(p_i(\boldsymbol{r}, \boldsymbol{d}(t_i)) || p_j(\boldsymbol{r}, \boldsymbol{d}(t_j))\right)$, the Kullback-Leibler divergence between the probability distributions for localization $i$ and $j$, which for 2 Normal distributions is defined as:

$$D_{\text{KL}}(i,j) = \frac{1}{2}\log\left(\frac{|\boldsymbol{\Sigma}_j|}{|\boldsymbol{\Sigma}_i|}\right) + \frac{1}{2}\text{Tr}\left(\boldsymbol{\Sigma}_j^{-1}\boldsymbol{\Sigma}_i\right) + \frac{1}{2}\left(\boldsymbol{\mu}_i - \boldsymbol{d}(t_i) - \boldsymbol{\mu}_j + \boldsymbol{d}(t_j)\right)^{\text{T}} \boldsymbol{\Sigma}_j^{-1}\left(\boldsymbol{\mu}_i - \boldsymbol{d}(t_i) - \boldsymbol{\mu}_j + \boldsymbol{d}(t_j)\right) - \frac{K}{2} \tag{S2}$$

The first term of Equation S3 can be dropped as it does not change under drift change. We define the optimization metric $G(\boldsymbol{D})$:

$$G(\boldsymbol{D}) = -\frac{1}{N}\sum_i^N \log\left(\sum_j^N \frac{1}{N} e^{-D_{\text{KL}}(i,j)}\right) \tag{S3}$$

The gradient of $G(\boldsymbol{D})$ w.r.t each drift parameter in $\boldsymbol{D}$ has to be computed in order to perform gradient descent. Taking the derivative it follows that:

$$\frac{\partial G(\boldsymbol{D})}{\partial \boldsymbol{D}} = -\sum_i^N \left(\sum_j^N e^{-D_{\text{KL}}(i,j)}\right)^{-1} \left(\sum_j^N \frac{\partial}{\partial \boldsymbol{D}} e^{-D_{\text{KL}}(i,j)}\right) \tag{S4}$$

$$\frac{\partial G(\boldsymbol{D})}{\partial \boldsymbol{D}} = \sum_i^N \left(\sum_j^N e^{-D_{\text{KL}}(i,j)}\right)^{-1} \left(\sum_j^N e^{-D_{\text{KL}}(i,j)} \frac{\partial}{\partial \boldsymbol{D}} D_{\text{KL}}(i,j)\right) \tag{S5}$$

To simplify, we define a normalization term:

$$Z_i(\boldsymbol{D}) = \sum_j^N e^{-D_{\text{KL}}(i,j)} \tag{S6}$$

$$\frac{\partial G(\boldsymbol{D})}{\partial \boldsymbol{D}} = \sum_i^N \sum_j^N \frac{1}{Z_i(\boldsymbol{D})} e^{-D_{\text{KL}}(i,j)} \frac{\partial}{\partial \boldsymbol{D}} D_{\text{KL}}(i,j) \tag{S7}$$

First, we derive the gradient of $G(\boldsymbol{D})$ w.r.t the drift in each frame. Recall the Kullback-Leibler divergence as defined in the main text (equation 5):

$$D_{\text{KL}}(i,j) = \frac{1}{2}\sum_{k=1}^K \left(\log\left(\frac{\sigma_{j,k}^2}{\sigma_{i,k}^2}\right) + \frac{\sigma_{i,k}^2}{\sigma_{j,k}^2} + \frac{\left(\mu_{i,k} - d_k(t_i) - \mu_{j,k} + d_k(t_j)\right)^2}{2\sigma_{j,k}^2}\right) - \frac{K}{2} \tag{S8}$$

Notice that the drift gradient of $D_{\text{KL}}(i,j)$ w.r.t. the drift of a frame $t$ is only nonzero if either localization $i$ or localization $j$ is part of frame $t$:

$$\frac{\partial}{\partial d_k(t)} D_{\text{KL}}(i,j) = \begin{cases} -\frac{1}{2\sigma_{j,k}^2}\left(\mu_{i,k} - d_k(t_i) - \mu_{j,k} + d_k(t_j)\right), & \text{if } t = t_i \\ \frac{1}{2\sigma_{j,k}^2}\left(\mu_{i,k} - d_k(t_i) - \mu_{j,k} + d_k(t_j)\right), & \text{if } t = t_j \\ 0, & \text{otherwise} \end{cases} \tag{S9}$$

The next goal is to collect all the gradient terms for a particular frame $t$:

$$\frac{\partial G(\boldsymbol{D})}{\partial \boldsymbol{d}(t)} = \sum_{i \in S_t} \sum_{j=1}^{N} \left( \frac{e^{-D_{\text{KL}}(i,j)}}{Z_i(\boldsymbol{D})} \frac{\partial}{\partial \boldsymbol{d}(t)} D_{\text{KL}}(i,j) \right) + \sum_{i=1}^{N} \sum_{j \in S_j} \left( \frac{e^{-D_{\text{KL}}(i,j)}}{Z_i(\boldsymbol{D})} \frac{\partial}{\partial \boldsymbol{d}(t)} D_{\text{KL}}(i,j) \right) \tag{S10}$$

Where $S_t = \{i : t_i = t\}$, the set of indices of localizations detected in frame $t$. We can combine these two sums into one by swapping $i$ and $j$ indices on the last term:

$$\frac{\partial G(\boldsymbol{D})}{\partial \boldsymbol{d}(t)} = \sum_{i \in S_t} \sum_{j=1}^{N} \left( \frac{e^{-D_{\text{KL}}(i,j)}}{Z_i(\boldsymbol{D})} \frac{\partial}{\partial \boldsymbol{d}(t)} D_{\text{KL}}(i,j) + \frac{e^{-D_{\text{KL}}(j,i)}}{Z_j(\boldsymbol{D})} \frac{\partial}{\partial \boldsymbol{d}(t)} D_{\text{KL}}(j,i) \right) \tag{S11}$$

For every localization, we collect a list of nearby localizations. The inner sum with index $j$ is evaluated fast because we can discard all the localizations $j$ that are far away from $i$. For those localizations, $e^{-D_{\text{KL}}(i,j)}$ will have dropped to effectively zero, removing any contribution to the gradients.

Finally, the gradient w.r.t spline control points ($\boldsymbol{u}_i$) can be computed using the chain rule:

$$\frac{d}{d\boldsymbol{u}_i} G(\boldsymbol{D}) = \sum_{t=1}^{L} \frac{\partial \boldsymbol{d}(t)}{\partial \boldsymbol{u}_i} \frac{\partial G(\boldsymbol{D})}{\partial \boldsymbol{d}(t)} \tag{S12}$$

#### 2. DRIFT ESTIMATION COMPUTE TIME

Figure S1 shows the compute time required for DME and RCC. For RCC, runtime is mostly dependent on the number of cross-correlation pairs that have to be computed ($n(n-1)/2$) where $n$ is the number of bins). The displayed compute time for DME includes 3 steps:

1. Initial estimate using RCC at 10 bins
2. DME estimate on the full dataset
3. DME estimates on the split datasets to estimate precision

It can be seen that DME takes around 1 minute or less, except when the binning size is set very small (1 or 2 frames).

#### 3. LOCALIZATION PIPELINE: CONVERTING RAW IMAGES TO LOCALIZATIONS

This section describes our custom localization pipeline, consisting of published work with some small details specific to our code. The processing involves the following steps:

- Raw camera pixel values stored as TIFF are converted into photon counts using calibrated camera gain and offset values. $N_{\text{photons}} = (I_{camera} - \text{Offset}) \cdot \text{Gain}$, where $I_{camera}$ is the light intensity on the pixel, in arbitrary ADC units from the camera.
- Spot detection and extracting small regions-of-interest (ROI) for later single molecule fitting.
- Maximum-likelihood fitting of the ROI pixels to a PSF model, resulting in 2D or 3D localizations. This is done differently depending on a 2D or 3D PSF model.
- Filtering the localizations based on the Cramer-Rao Lower Bound (CRLB), or $\chi^2$ metric.

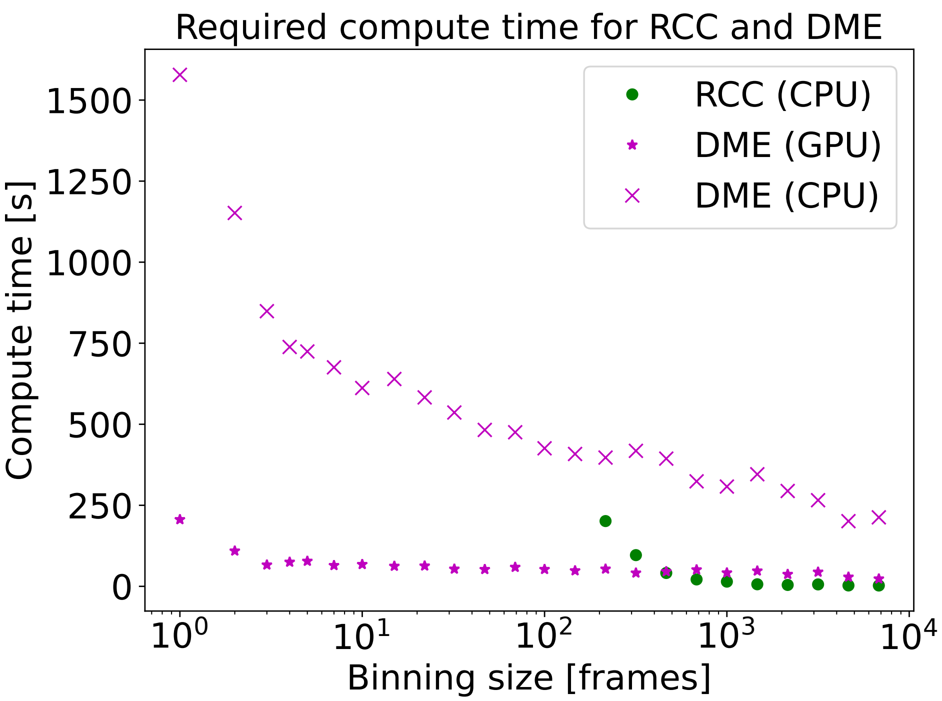


**Fig. S1.** Run time of the 2 drift estimation algorithms, measured on an Alienware m15 R3 laptop with i9-10980 CPU and nVidia Geforce RTX 2080 GPU. The dataset is from SIMFLUX [2], as displayed in Figure 4 of the main text, with 25000 frames and 820802 localizations. Missing RCC values indicate binning sizes where there are too few localizations in a particular bin and Picasso report "RCC failed".

##### A. Spot detection

To allow for effective 3D spot detection, we use a template based spot detection scheme. The template is a Z-stack of the PSF at various positions relative to the focal plane. For every Z position, we compute the 2D correlation with the PSF at that position using fast fourier transforms:

$$A_z = \mathcal{F}^{-1}\{\mathcal{F}\{I - Bg\} \cdot \mathcal{F}\{PSF_z\}\} \tag{S13}$$

Here, $Bg$ is either a static background image or a dynamic estimate of the background, such as compute the recorded movie filtered by a temporal median filter. 2D spot detection is typically implemented using a series of 2D uniform and maximum filters[3]. Similarly, we apply a 3D max-filter is applied to the resulting Z stack $A = [A_1 \cdots A_m]$:

$$A_{max} = max[A, L_{xy}, L_z] \tag{S14}$$

Here, $A_{max}$ is the output of a 3D maximum filter applied on the stack of PSF correlations $A$, with a window size of $L_{xy} = 5\sigma_{PSF}$ and $L_z = m$ (over the full Z range). $\sigma_{PSF}$ indicates the approximate width of a Gaussian-like PSF. The value is manually tuned for other kinds of PSF shapes.

Finally, $A_{max}$ is compared to $A$ to find peaks:

$$A_{peak} = \begin{cases} A, & \text{if } A_{max} = A \\ 0, & \text{otherwise} \end{cases} \tag{S15}$$

Spots are detected whenever $A_{peak}$ is higher than a preset tuned threshold value, and a region-of-interest is extracted around that peak pixel.

##### B. 2D Localization

Fitting of 2D localization data from is done using the Levenberg-Marquardt algorithm for maximum-likelihood estimation on Poisson distributed samples [4]. The model is a 2D Gaussian PSF modelS16 that integrates the 2D Gaussian over the pixel area[5].

$$\mu_k = \theta_{bg} + \theta_I \frac{1}{2\pi\sigma_x\sigma_y} \iint_{A_k} \exp\left(-\frac{(x-\theta_x)^2}{2\sigma_x^2} - \frac{(y-\theta_y)^2}{2\sigma_y^2}\right) dxdy \tag{S16}$$

Here, $\mu_k$ is the expected value for pixel $k$. The fitted parameters are as follows: $\theta = \{\theta_x, \theta_y, \theta_I, \theta_{bg}, \theta_{sigma_x}, \theta_{sigma_y}\}$. $A_k$ is the area of pixel $k$.

We noticed that fitting more parameters including the Gaussian width ($\sigma_x$) and height ($\sigma_y$) will lower precision and make the estimation less robust. For this reason the fitting is split up in 3 different steps:

1. Fit all detected spots using a fixed-sigma PSF model that includes $\theta_x$, $\theta_y$, $\theta_I$, $\theta_{bg}$ (background photons/pixel) as fitted parameters. This initial sigma value is a value entered by the user.
2. Using the previous positions as initial estimates, re-fit the data now with a model that includes Gaussian $\sigma_x$ and $\sigma_y$.
3. Bin all localizations over time bins, and compute the median $\sigma_x$ and $\sigma_y$ (see figure S2). These spline-interpolated PSF sigma's are now used for a final localization run, improving the localization precision and robustness. This effectively corrects for Z drift without requiring a 3D PSF model.

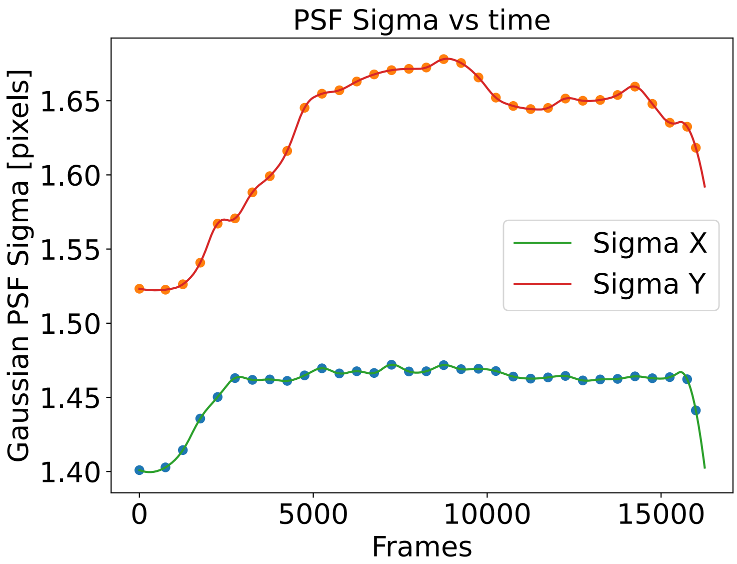


**Fig. S2.** For 2D localization such as used in total internal reflection (TIRF) microscopy, typically a 2D Gaussian PSF model is used. In our pipeline, we bin all fitted Gaussian sigma values, compute the median values for each bin, and interpolate these between time bins. The data shown above are PSF parameters for the DNA-PAINT measurement shown in figure 2 in the main text. Each data point is the median $\sigma_x$ and $\sigma_y$ value for a bin of 500 frames.

##### C. 3D Spline-based localization

Fitting of 3D data is done using a 3D cubic spline model. First, the PSF is estimated from bead data using MATLAB code[6]. Then, we use the resulting .MAT file in our custom written localization pipeline to perform localization, again using the Levenberg-Marquardt algorithm [4]. For 3D localization, we find that convergence of the MLE is best when the parameters are updated using the scale invariant approach [7]

##### D. Python package

The DME algorithm is integrated into our Python/C++/CUDA localization pipeline called "photonpy", but can also be build independently from source.

Photonpy localization pipeline:
https://github.com/qnano/photonpy

Drift estimation example code:
https://github.com/qnano/drift-estimation
